# Interoceptive training enhances emotional awareness and body image perception: evidence from improved heartbeat detection and psychological outcomes

**DOI:** 10.1101/2025.06.16.659873

**Authors:** Anna Rusinova, Vladislav Aksiotis, Ekaterina Potapkina, Elizaveta Kozhanova, Vladislav Akimov, Alexei Ossadtchi, Maria Volodina

## Abstract

The conscious integration of interoceptive signals, such as heartbeats and breathing, is essential for emotional regulation, self-awareness, and stress resilience. We conducted a controlled trial implementing a 5-day biofeedback-guided interoceptive training program to assess its impact on interoceptive accuracy (IAc) and psychological dimensions of bodily and mental perception. The experimental group showed significant improvements in IAc compared to controls, reflecting enhanced detection of heartbeats. Training increased scores on two MAIA subscales: “Awareness of Negative Emotional States” and “Not Distracting”. These outcomes suggest the intervention enhanced participants’ capacity to acknowledge and engage with unpleasant bodily sensations without resorting to avoidance or distraction strategies. A trend toward improved body image perception emerged in the experimental group, with gains correlating with reductions in depressive symptoms, underscoring the interplay between bodily awareness and emotional well-being. These findings underscore that targeted interoceptive training represents a powerful tool for refining awareness of internal states, enhancing attentional control, and optimizing emotional processing. By bridging bodily signals and affective experiences, this approach offers a pathway for innovative, evidence-based therapies to bolster emotional resilience and address conditions rooted in dysregulated emotional states, such as anxiety, depression, and stress-related disorders.

## Introduction

Interoception, the process of perceiving and interpreting internal bodily signals such as heartbeat, respiration, and hunger, plays a crucial role in emotion regulation ^1, 2^. As a core mechanism of self-perception and subjective experience, interoception enables individuals to monitor their physiological states and adaptively respond to emotional challenges ^3^.

Interoception emerges from the integration of multiple sensory inputs, including both physiological and subjective experiences ^4^. It encompasses autonomic, endocrine, and immune processes, supporting memory, affective and emotional experiences, and the psychological sense of self, with the insular cortex serving as a critical interface between the brain and body ^5^. This ability to monitor and interpret internal physiological states is essential for maintaining homeostasis, driving motivation, and regulating autonomic, cognitive, and behavioral processes ^6^. Crucially, interoceptive processing is intricately linked to emotion generation, shaping both adaptive and maladaptive emotional responses.

Empirical evidence underscores the intimate connection between interoception and emotional processing. The somatic marker hypothesis posits that emotional processes are integral to decision-making, as they rely on bodily sensations associated with emotions ^7^. Empirical evidence supports this notion, showing that individuals with greater interoceptive accuracy (IAc), the ability to precisely detect and discriminate internal bodily signals, often assessed through tasks like heartbeat perception ^8^ and bodily awareness experience more intense emotional reactions ^9^ and exhibit heightened sensitivity to the emotions of others ^10^. Furthermore, interoception has been shown to be positively correlated with emotion regulation and stress resilience, with reduced IAc observed following acute stress ^11^. IAc are associated with more effective regulation of negative emotions and better adaptation to social uncertainty ^12^. Interoceptive awareness, which refers to the conscious recognition and interpretation of these internal signals, facilitates somatic reappraisal, integrating bodily sensations with explicit narratives to promote adaptive emotional processing ^13^.

Disruptions in interoceptive processing, a critical mechanism underlying emotional experience and regulation, contribute to the pathophysiology of diverse physical and mental disorders ^14^. These include anxiety ^15^, obsessive-compulsive disorder (OCD) ^16^, post-traumatic stress disorder ^17^, anorexia ^18^, bulimia ^19^, schizophrenia ^20^, functional motor disorder ^21^, and neurodegenerative conditions ^22^. Interoception is critically involved in both the emergence and regulation of affective states and is frequently altered in clinical and neurodevelopmental disorders ^23^. The mechanisms linking interoceptive dysfunction to psychopathology involve disruptions in neural and cognitive processes. For instance, aberrant fear-specific cardiac responses ^24^ and impaired integration of interoceptive signals within insular-frontotemporal networks ^25^ undermine the brain’s ability to contextualize bodily states, exacerbating affective dysregulation. Such deficits also hinder emotional awareness: individuals with poor IAc struggle to verbalize emotions ^1^ and mitigate the impact of negative experiences, as seen in depression ^26^. Beyond psychopathology, impaired IAc correlates with non-planning impulsivity ^27^ and poorer exercise tolerance in cardiac rehabilitation ^28^, underscoring its broad relevance to adaptive functioning.

Given the critical role of interoception in emotional regulation, interventions aimed at enhancing interoceptive awareness have gained prominence. Mindfulness-based practices, yoga, and biofeedback training have shown promise in improving internal body awareness and physiological regulation, leading to greater emotional control ^29, 30, 31^. Body and mental awareness training has gained recognition as a viable approach for enhancing emotional regulation and self-awareness. For example, a seven-week interoception-focused program significantly improved emotional regulation in children with diverse diagnoses in a special education setting ^30^. Similarly, Mindful Awareness in Body-oriented Therapy (MABT), has been shown to enhance interoceptive awareness, emotional regulation, and treatment outcomes in women with substance use disorders ^31^. However, the effectiveness of these interventions varies across populations, emphasizing the need for tailored approaches. Notably, both subjective interoceptive awareness and objective IAc appear to be modifiable through targeted interventions. The single yoga session enhanced IAc in healthy individuals but failed to do so in patients with anorexia nervosa, potentially due to deficits in emotional awareness and bodily signal processing ^16^. Moreover, research has demonstrated that self-focused attention tasks can have differential effects on IAc, enhancing it in healthy individuals while paradoxically decreasing it in those with anorexia ^32^. However, more sustained interventions appear to be effective; an eight-session yoga program successfully improved IAc in individuals with eating disorders ^33^.

Given the critical role of interoception in emotional regulation, approaches aimed at enhancing interoceptive functioning could be particularly promising for improving emotion regulation. This is especially important as existing methods, such as pharmacological treatments and cognitive-behavioral interventions, have several limitations. While pharmacotherapy effectively alleviates symptoms, it often lacks individualization ^34^ and may fail to address the underlying physiological dysregulation. Moreover, pharmacological treatments can lead to adverse effects such as emotional blunting, apathy, and a reduced ability to express emotions, including the inability to cry ^35^. In some cases, cognitive-based interventions have been found to be more effective than pharmacotherapy in regulating emotions ^36^. However, cognitive-behavioral therapy (CBT) also has limitations, including modest effect sizes and limited implementation in clinical settings ^37^.

In light of these limitations, interventions targeting body awareness and interoception have emerged as promising alternatives for enhancing psychophysiological well-being and emotional regulation. Such practices have been shown to significantly reduce stress and anxiety levels ^38^, enhance self-confidence ^39^, and improve IAc ^40, 41^. However, the effects of mindfulness-based interventions remain inconsistent, as some studies report no significant improvements in interoceptive or emotional awareness ^42^.

Interventions specifically designed to target interoception have demonstrated notable clinical efficacy across various treatment settings ^43^. Incorporating interoceptive awareness assessment into clinical practice may enhance the understanding of mental disorders, optimize patient care, and refine therapeutic strategies ^44^. Among these approaches, biofeedback-based training has shown particular promise by providing real-time physiological feedback, enabling individuals to regulate their autonomic responses more effectively. For example, contingent cardiac feedback training significantly increases IAc compared to non-contingent feedback, mindfulness practice, or passive waiting ^45^. However, findings on biofeedback interventions remain mixed, with some studies reporting no significant improvements in IAc ^46^, while others suggest that heart rate variability biofeedback therapy shows mixed results in improving interoception, with stronger effects when an intense treatment protocol is used ^47^.

Our study seeks to develop and evaluate a novel interoceptive training protocol aimed at enhancing awareness of physiological states and emotional experiences. Specifically, we hypothesize that biofeedback-based interoceptive training, supplemented with brief self-focused attention exercises, will not only improve interoceptive accuracy (IAc) but also foster positive changes in psychological factors and body awareness, as assessed through validated questionnaires. The primary goal of this research is to systematically investigate the effects of interoceptive training on both physiological and emotional processes. By doing so, we aim to provide valuable insights into innovative preventive strategies for mitigating psychophysiological distress and promoting overall well-being. This work may pave the way for more personalized and effective interventions targeting emotional health and bodily awareness.

## Results

Prior to the intervention, statistical analysis was conducted to assess baseline differences between the experimental and control groups. The Mann-Whitney U test revealed no significant differences in age (U = 191.5, p-value = 0.75) or baseline scores on interoceptive accuracy (IAc) (U = 197, p-value = 0.95) (See Supplementary material). Categorical variables were analyzed using the chi-square test of independence, which showed no significant differences in sex distribution (χ^2^(1) = 0.00, p-value = 1.00), education level (χ^2^(3) = 1.05, p-value = 0.79), relationship (χ^2^(2) = 0.36, p-value = 0.83) or family status (χ^2^(3) = 4.53, p-value = 0.21).

### Interoceptive accuracy

A Mixed Linear Model Regression analysis was conducted to evaluate changes in IAc after intervention. IAc served as the dependent variable, with fixed effects including time point (before/after) and group (experimental/control). The analysis revealed a significant interaction between group and time point (p-value = 0.001), indicating that the trajectories of IAc changes differed between the experimental and control groups. Additionally, the Wilcoxon matched pairs test confirmed a significant increase in IAc within the experimental group (before: 0.75 (IQR 0.59 - 0.79), after: 0.84 (IQR 0.76 - 0.87), p-value < 0.01). While no significant changes were observed in the control group (before: 0.7 (IQR 0.58 - 0.86), after: 0.73 (IQR 0.67 - 0.84), p-value > 0.1) (See Fig 3.). These findings indicate that interoceptive training effectively enhances participants’ ability to accurately perceive internal bodily signals.

**Fig 1.**
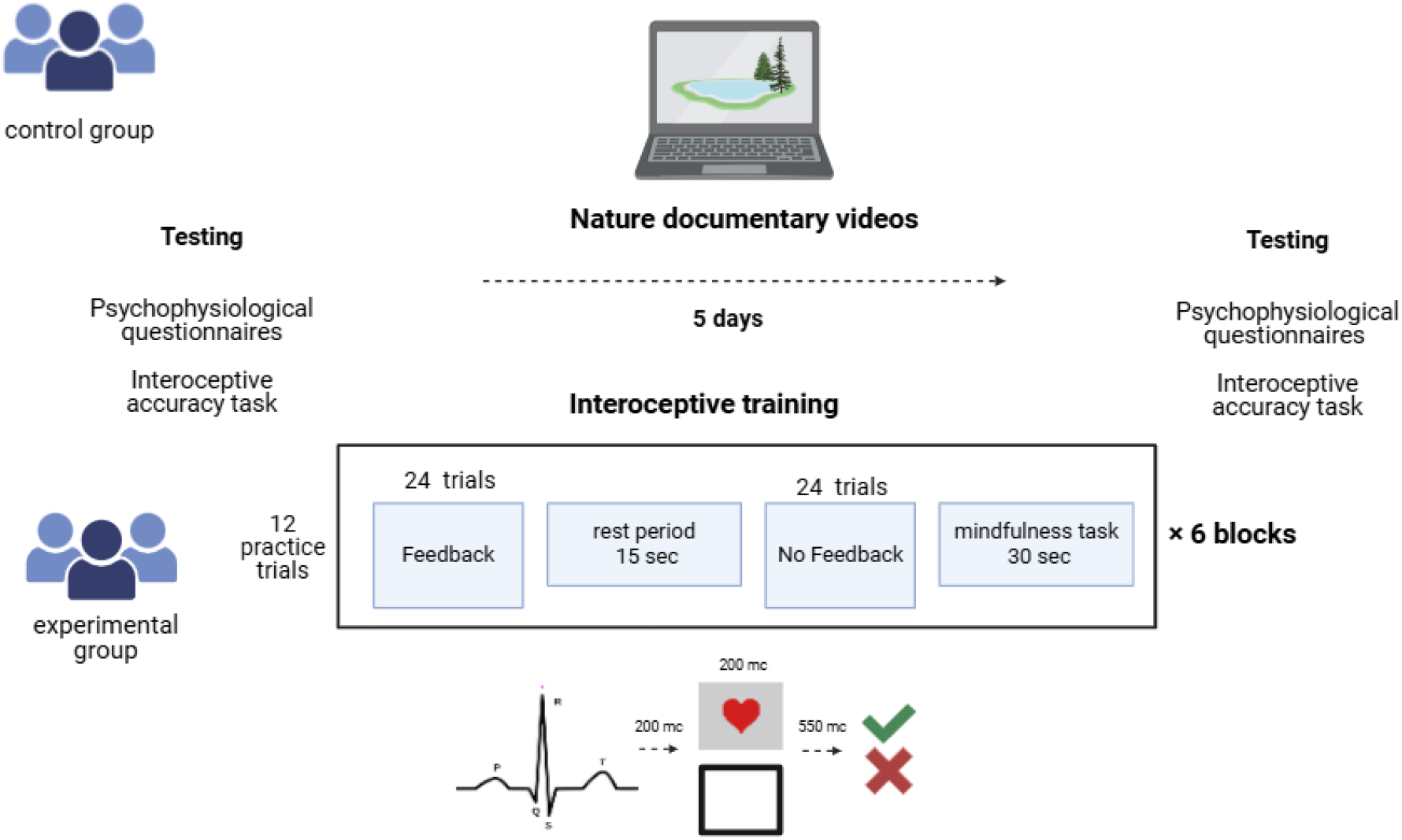
The design of the experiment.

**Fig 2.**
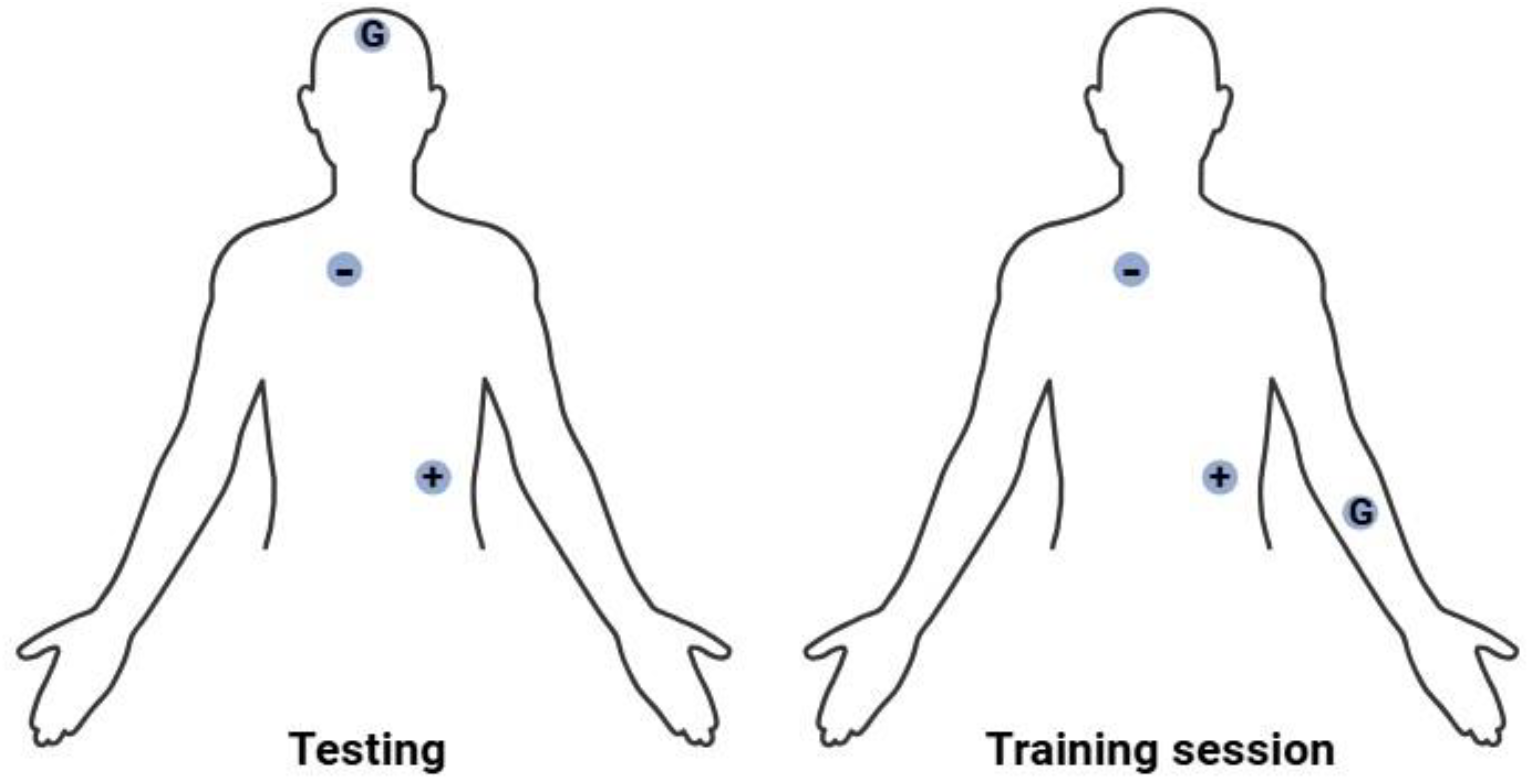
ECG electrodes placement

**Fig 3.**
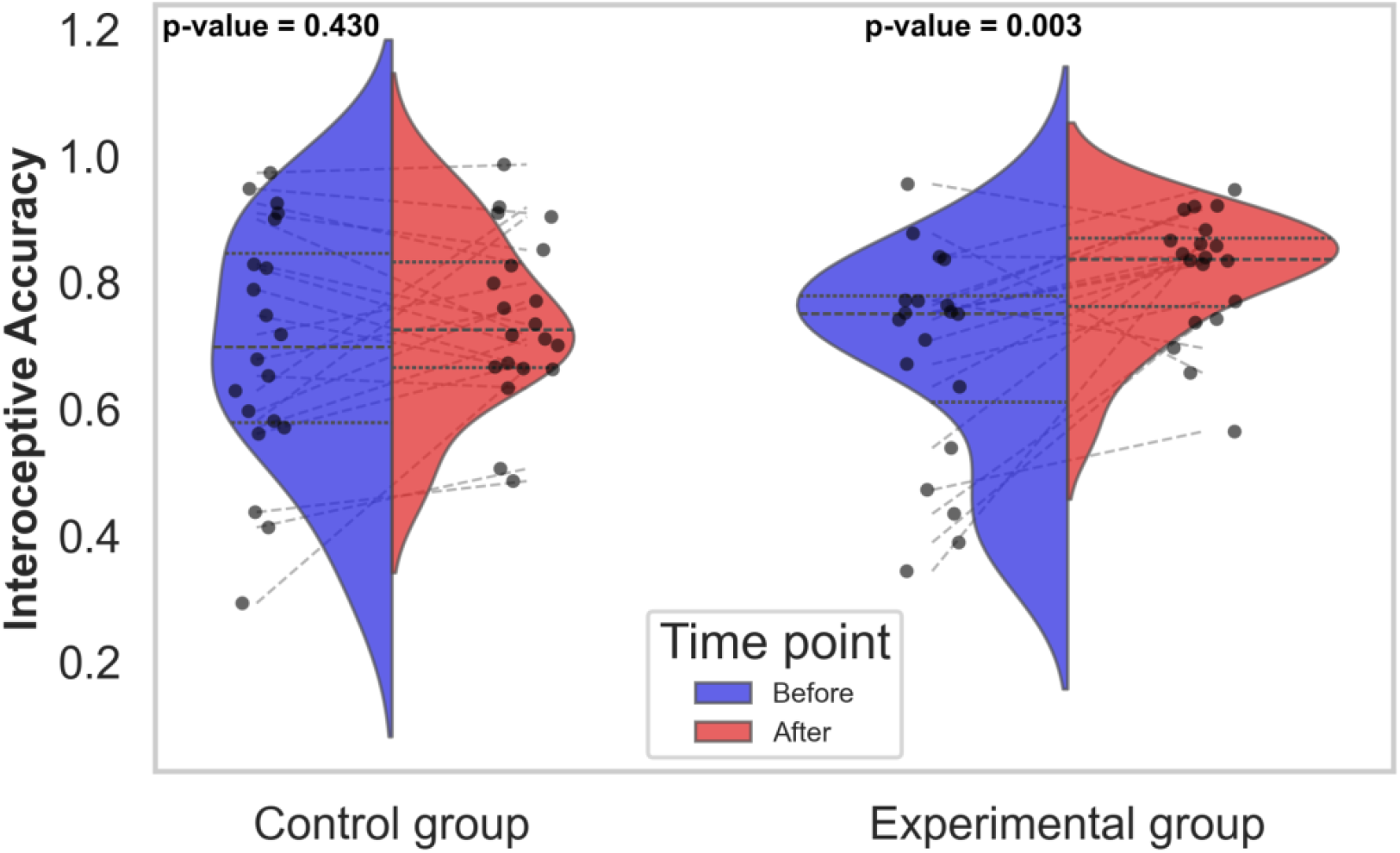
Violin plot illustrating the distribution of IAc before and after training in the control and experimental groups. The blue area represents IAc distribution before the intervention, the red area represents IAc distribution after the intervention. Black dots indicate individual data points, and gray lines connect repeated measures within the same participant. The central dark line within each violin represents the median (50th percentile). The upper and lower dashed lines correspond to the 75th and 25th percentiles, respectively, marking the interquartile range (IQR). The p-values were obtained using the Wilcoxon signed-rank test.

### Questionnaires

The Mixed Linear Model Regression analysis revealed significant group-by-time interactions for the MAIA (Multidimensional Assessment of Interoceptive Awareness) subscales “emotional awareness of negative state” (p-value = 0.042) and “not distracting” (p-value = 0.047), indicating that the training had a differential impact on these aspects of interoceptive awareness. The Wilcoxon matched pairs test further confirmed a significant improvement in the experimental group for both “not distracting” (before: 1.33 (IQR 0.67 - 2.33), after: 2.33 (IQR 1.67 - 3.0), p-value < 0.01) and “emotional awareness of negative state” (before: 2.33 (IQR 1.67 - 3.67), after: 3.0 (IQR 2.17 - 4.0), p-value < 0.02), while no significant changes were observed in the control group (p-value > 0.5) (See Fig 4.). This demonstrates that interoceptive training enhances participants’ ability to remain attentive to unpleasant bodily sensations rather than ignoring or distracting themselves from them. Additionally, it improves their capacity to recognize and process negative emotional states effectively.

**Fig 4.**
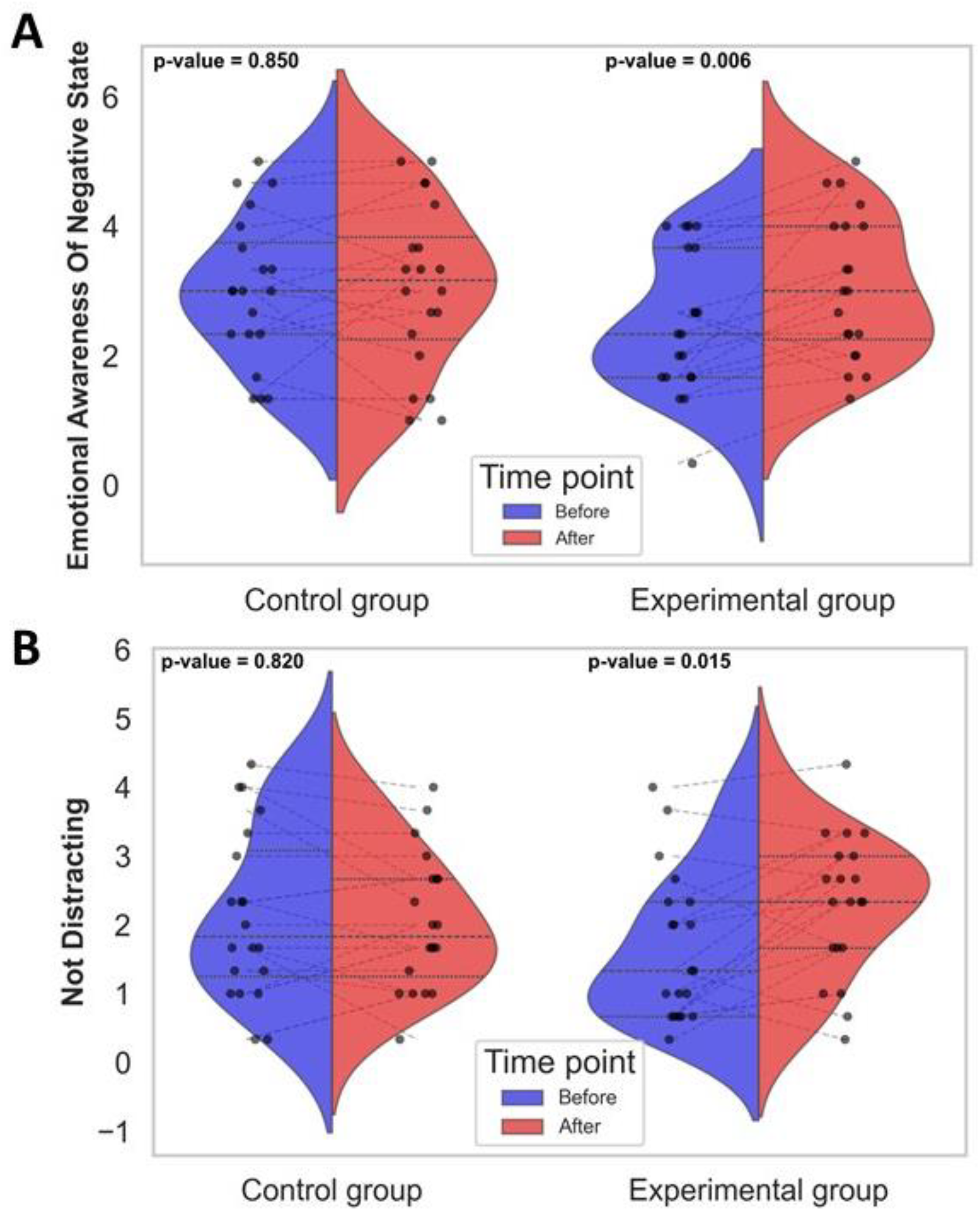
Violin plot illustrating the distribution of “emotional awareness of negative state” (A) and “not distracting” (B) scores before and after training in the control and experimental groups. The blue area represents scores distribution before the intervention, the red area represents scores distribution after the intervention. Black dots indicate individual data points, and gray lines connect repeated measures within the same participant. The central dark line within each violin represents the median (50th percentile). The upper and lower dashed lines correspond to the 75th and 25th percentiles, respectively, marking the interquartile range (IQR). The p-values were obtained using the Wilcoxon signed-rank test.

**Fig 5.**
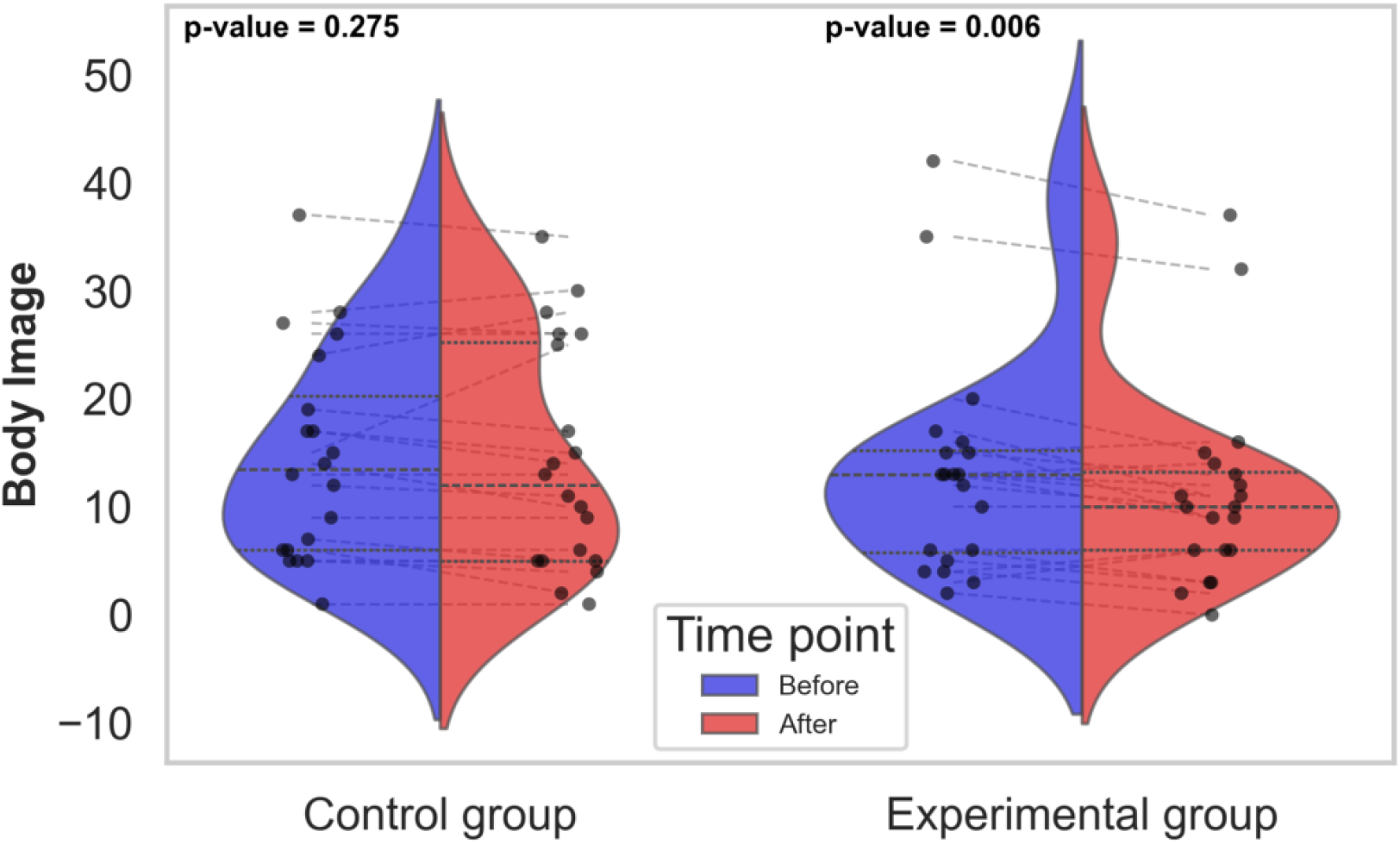
Violin plot illustrating the distribution of body image scores before and after training in the control and experimental groups. The blue area represents scores distribution before the intervention, the red area represents scores distribution after the intervention. Black dots indicate individual data points, and gray lines connect repeated measures within the same participant. The central dark line within each violin represents the median (50th percentile). The upper and lower dashed lines correspond to the 75th and 25th percentiles, respectively, marking the interquartile range (IQR). The p-values were obtained using the Wilcoxon signed-rank test.

The Mixed Linear Model Regression analysis revealed a trend toward significance in the group-by-time interaction for body image scores (p-value = 0.068). The Wilcoxon matched pairs test further confirmed a significant decrease of body image scores in the experimental group (before: 13.0 (IQR 5.5 - 15.5), after: 10.0 (IQR 6.0 - 13.5), p-value < 0.01), indicating a reduction in body concerns. Whereas no significant changes were observed in the control group (p-value > 0.05), see Fig 4. These results indicate that interoceptive training may contribute to a more positive body image perception.

For other psychological measures, no significant interaction was found between group and time point factors.

### Correlation analysis

For the variables that demonstrated statistically significant changes in the Wilcoxon matched pairs test, we conducted correlation analyses to explore the relationships between these changes and alterations in other indicators. Spearman’s rank correlation analysis revealed significant associations within the experimental group, indicating that improvements in body image perception were positively correlated with a reduction in depressive symptoms (r = 0.65, p < 0.01) following the training.

## Discussion

The results of our study indicate that a short course of interoceptive training, lasting only five days, can enhance interoceptive accuracy in the Heartbeat Counting Test and influence specific dimensions of interoceptive awareness associated with emotion regulation skills. The training protocol we developed combined ECG-based biofeedback training with mindfulness techniques. Both approaches have been previously shown to be effective in enhancing interoceptive sensitivity. For instance, eight biofeedback sessions over five weeks (1–3 per week) led to a significant increase in IAc ^41^, while in our protocol, comparable improvements were achieved within just five sessions over one week. Similarly, a study with three heartbeat perception biofeedback sessions found significant improvements only after the first 20-minute session, with no further gains over three weeks ^48^, possibly due to a learning effect from repeated IAc assessments. Another study showed that a single 1.5-hour biofeedback session enhanced heart rate discrimination when using tactile, but not visual, feedback ^49^, whereas in our protocol, visual feedback alone was sufficient to improve IAc. While biofeedback-based interventions have demonstrated efficacy, other body-oriented approaches, such as mindfulness and meditation, have shown mixed effects. A body scan intervention (20 minutes daily for eight weeks) significantly improved IAc ^39^, and a nine-month contemplative mental training program (13 weekly 2-hour group sessions plus 30 minutes of daily practice) led to improvements emerging after six months ^40^. We propose that integrating these two methods may be a promising strategy for increasing training efficacy and reducing the duration required to achieve noticeable effects.

It is important to note that an increase in interoceptive accuracy can be associated with both positive effects on an individual’s psycho-emotional state ^9^, and heightened levels of anxiety ^50^. Therefore, to discuss the overall positive impact of interoceptive training, it is crucial to evaluate its effects on the psychological well-being of participants. The ability to accurately perceive and interpret interoceptive cues plays a crucial role in emotional regulation, stress resilience, and mental health ^51^. Research suggests that deficits in interoception are linked to various psychological disorders, including anxiety ^52^, depression ^53^, and alexithymia ^54^. It is assumed that in clinical populations, the effects of interoceptive interventions may be more complex and context-dependent ^55^, while studies have largely reported positive outcomes. For instance, improving heart rate perception in patients with somatoform disorders has been shown to significantly reduce the severity of distress symptoms ^56^. A seven-week intervention led to better emotional regulation in children with various diagnoses from a special educational classroom ^30^. Additionally, interoceptive interventions are hypothesized to be particularly effective in reducing physiological symptoms associated with autonomic dysregulation ^57^. However, not all studies have demonstrated consistent benefits across different populations. For example, while a single 75-minute hatha yoga session improved interoceptive accuracy in healthy individuals, it did not yield the same effect in individuals with anorexia nervosa ^16^, suggesting that clinical impairments may limit learning effects observed in healthy participants. An important direction for future research is the integration of interoceptive training with other interventions, such as cognitive-behavioral therapy ^58^. By developing targeted training protocols, individuals may enhance their ability to recognize and interpret physiological signals, thereby promoting adaptive emotional responses and mitigating maladaptive stress reactions. Moreover, such interventions have the potential to improve self-awareness and mindfulness, contributing to greater psychological stability and overall well-being. It is important to note, however, that our study was conducted exclusively on healthy participants. Determining the most effective training protocols and durations for specific clinical populations remains a critical area for further investigation.

Our results revealed significant improvements in the MAIA subscales “not distracting” and “emotional awareness of negative state” following the training, along with a notable trend toward decreased “body image” scores with no changes in depression, anxiety and emotionality scores.

The “emotional awareness of negative state” subscale in the MAIA (Multidimensional Assessment of Interoceptive Awareness) questionnaire assesses an individual’s ability to recognize and acknowledge their own negative emotional states, as well as the associated bodily sensations. This subscale reflects how aware a person is of the physical manifestations of emotions like anxiety, sadness, or anger and whether they can identify these feelings without ignoring or suppressing them ^59^. In the study by Mehling et al. it was found that “emotional awareness”, including “emotional awareness of negative state” exhibited a negative correlation with trait anxiety when analyzed in isolation. However, this relationship shifted to a positive correlation once the variance shared with “self-regulation” was accounted for. Mehling et al. concluded that simply being aware of how bodily sensations relate to emotional states without the capacity to utilize that awareness for distress reduction could paradoxically heighten anxiety levels. This underscores that enhancing interoceptive sensibility is not a universal solution; it necessitates a nuanced understanding of the regulatory and attentional processes embedded within the construct of interoceptive sensibility, which holds significant implications for clinical practice.

Considering this, it is essential to highlight that the training did not result in heightened levels of anxiety or emotional reactivity. In contrast, participants demonstrated improved scores in the “not distracting” category. The “not distracting” subscale in the MAIA questionnaire evaluates an individual’s ability to not to ignore or distract themselves from sensations of pain or discomfort. This dimension reflects how well a person can remain aware of and acknowledge unpleasant bodily sensations without resorting to avoidance or distraction strategies ^60^.

The ability to not distract oneself from negative sensations represents a promising strategy in managing chronic pain ^61^. This approach, based on mindful awareness of one’s bodily state, serves as an alternative to coping strategies that rely on distraction and ignoring pain. By deepening awareness of their sensations, individuals may improve pain management and overall well-being.

In clinical settings, an increased focus on physical sensations is often linked to anxiety, hypervigilance, somatization, and hypochondriasis, leading to a perception of maladaptive interoceptive awareness. However, recent research has highlighted an alternative approach, mindful rather than anxiety driven, which offers positive effects and enhances emotional regulation ^60^. This shift is supported by studies demonstrating that trusting bodily signals and maintaining attention on internal states are essential for emotional regulation ^62^.

Additionally, interoceptive training resulted in a decrease in body image scores in the experimental group, indicating a reduction in body-related concerns. This outcome is consistent with the idea that improving interoceptive awareness may be beneficial for body image perception. Previous research has suggested that interoceptive training may be beneficial for body image perception, as disruptions in interoceptive processing are closely associated with eating disorders, including anorexia nervosa ^18^ and bulimia nervosa ^19^, it can be hypothesized that enhancing interoceptive abilities might contribute to improved body image perception and related mental health outcomes. These findings align with existing literature suggesting that interoceptive awareness plays a key role in body satisfaction and emotional well-being ^63^. A possible mechanism underlying this relationship is increased somatic self-awareness, which fosters a more attuned and accepting perception of one’s body, thereby mitigating negative self-perceptions and depressive symptomatology. Beyond its implications for eating disorders, interoception-based interventions have shown promise in improving body image across diverse populations. For instance, an embodied rehabilitative training program significantly enhanced interoceptive awareness, body image perception, balance, and quality of life in patients with mild multiple sclerosis ^64^.

## Conclusion

Our study demonstrates the efficacy of a brief, five-day interoceptive training protocol that integrates biofeedback and mindfulness practices to enhance interoceptive accuracy while positively influencing psychological well-being. Notably, our findings suggest that even a short-term intervention can yield dual benefits: improving physiological awareness and potentially fostering emotional regulation. Given the established link between disrupted interoceptive processing and conditions such as body image concerns, anorexia nervosa, and bulimia nervosa, this protocol holds promise for addressing both physical and emotional health challenges.

The ability to cultivate interoceptive awareness within a concise, one-week framework is particularly valuable for practical applications. This protocol can be seamlessly integrated into hospitalization or rehabilitation programs, providing accessible support for individuals with limited time availability, including athletes, professionals, and clinical populations requiring efficient interventions. Furthermore, the observed improvements in interoceptive accuracy and psychological outcomes highlight its potential for enhancing emotional regulation and stress management.

Our findings contribute to the growing body of research on interoceptive training and its implications for mental health, underscoring its relevance in self-regulation and therapeutic contexts ^29, 65^. Future research should explore individual differences in responsiveness to interoceptive training, refine optimal protocols, and assess the long-term effects on mental health, particularly in populations prone to emotional dysregulation or low interoceptive sensitivity. By integrating neurobiological and psychological perspectives, we can deepen our understanding of the mechanisms underlying interoceptive training and expand its application in promoting holistic well-being.

## Methods

### Participants

The study included 40 participants. The inclusion criteria were as follows: participants were selected from individuals aged 18 to 40 years, participants should be without diagnosed mental illnesses or brain disorders and not taking drugs that affect the central nervous system, such as antidepressants or sedatives. One participant was excluded from the final analysis because his/her IAc scores failed to meet the criteria of the statistical test for outliers. Thus, the data from 39 individuals were included in the final analysis with thirteen men and twenty-six women (mean age = 22.73, SD = 5.39) for IAc analysis. The experiment was conducted in accordance with the Declaration of Helsinki and approved by the HSE University Committee on Inter-University Surveys and Ethical Assessment of Empirical Research (#3 from 03.04.2024). Participants received compensation (20 USD at purchasing power parity). All participants provided written informed consent.

### Procedure

The design of the experiment is shown in Figure 1.

Before the experimental session, all participants completed a series of psychological questionnaires (see below). Then, participants were instructed to sit comfortably and relax for 3 minutes, after which they were given a heartbeat tracking task ^66^ designed to assess IAc ^67^. IAc was evaluated according to the following formula:

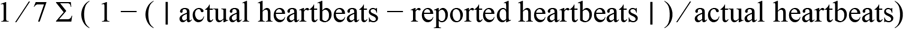

Before each counting interval, participants were prompted with the message “Get ready” displayed on the screen for 3.5 seconds. During this time, they were instructed to place their hands comfortably and prepare to count their heartbeats. Participants were then asked to silently count their heartbeats over specific durations of 20, 25, 35, 45, 55, 65, 75 seconds without physically measuring their pulse or using any other methods to detect their heartbeats, while “Counting” was on the screen.

At the end of each counting interval, participants reported the number of heartbeats they had counted. They did not receive any feedback about the accuracy of their counts or the duration of the intervals. As highlighted in prior research ^68, 69^, explicit knowledge of heart rate can distort performance on interoceptive tasks. To address this, participants in our study were not informed of the interval durations, thereby eliminating the potential for time-based estimation of heart rate.

### Intervention

The experimental group completed a 5-day interoceptive training protocol, while the control group watched nature documentary videos for 40 minutes each day over the same period. Each session of interoceptive training lasted about 40 minutes.

Interoceptive training consisted of five sessions conducted daily. Participants were instructed to press a button after a specific number of heartbeats (2, 3, or 4), as indicated on the screen at the beginning of each trial. The training session was divided into six distinct blocks. Each block was divided into two stages. In the first stage, a heart image was displayed for 200 milliseconds after every R-peak detection. In the next stage, no visual feedback was provided. Participants were required to rely solely on their internal bodily sensations to monitor their heartbeats and press the button after the instructed number of beats. A response was considered correct if the button press occurred within 0.55 seconds following the heartbeat detection, considering a physiological delay of 200 milliseconds. Visual feedback was provided immediately after each trial: a green check mark indicated a correct response, while a red X sign indicated an incorrect response. After every block participants engaged in brief mindfulness-based focus periods, concentrating on their bodily sensations and practicing deep, slow breathing for 30 seconds. After each stage in the block, a 15-second rest period was provided to ensure participants could reset and prepare for the next stage. The whole session lasted approximately forty minutes. Stimuli were presented and responses were recorded using PsychoPy ^70^.

Psychological questionnaires were administered to participants in the laboratory prior to physiological testing, before the attachment of all equipment, and again after the intervention following the removal of all equipment. Since the first and final intervention sessions took place on the same days as questionnaire completion and physiological testing, there was an approximately one-hour resting interval between the intervention and subsequent testing.

### ECG recording

Throughout the experiment, ECG was recorded using an NVX52 amplifier (Medical Computer Systems, Russia), NeoRec software (Medical Computer Systems, Russia) and digitized at a sampling rate of 500 Hz. The first ECG electrode was positioned inferior to the right clavicle, while the second electrode was placed along the left midaxillary line (Figure 2). This configuration was selected to maximize signal quality and minimize motion artifacts during data acquisition. During testing, the ground electrode was placed on the head to facilitate synchronization with other physiological recordings, such as EEG. Synchronization of the ECG signal and stimulus presentation was performed using a photo sensor fixed in the upper right corner of the monitor.

### ECG data processing

#### ECG data processing in IAc task

Synchronization between ECG signal acquisition and stimulus presentation was achieved using a photo sensor attached to the upper right corner of the monitor, ensuring precise alignment of physiological data and task events. Before the start of the interoceptive accuracy assessment, a 3-minute resting baseline recording was conducted to establish each participant’s individual heart rate variability. During the task, the recorded ECG signals underwent preprocessing, including a 0.5 Hz high-pass filter to eliminate slow signal drifts, a 70 Hz low-pass Butterworth filter to remove high-frequency noise, and a 50 Hz notch filter to suppress power line interference. R-peaks were detected using the bio_process function from the NeuroKit2 library ^71^.

Event markers corresponding to the start of each trial were extracted using the MNE-Python library, with onset times aligned to stimulus presentation. A bandpass filter from 5 to 50 Hz was applied to focus on the relevant physiological frequency band for accurate R-peak detection. During the task, real-time detection of R-peaks initiated the delivery of feedback stimuli.

### ECG data processing in training session

During the training session, ECG data were recorded at a sampling rate of 250 Hz. The signal was transmitted using the Lab Streaming Layer (LSL) protocol, enabling real-time synchronization between physiological measurements and the visual feedback provided to participants.

To ensure accurate detection of R-peaks and minimize signal noise, the raw ECG data were preprocessed using a series of filters. A 5 Hz high-pass filter was applied to remove slow drifts, a 50 Hz low-pass Butterworth filter eliminated high-frequency noise, and a 50 Hz notch filter reduced interference from power line noise.

R-peak detection was performed using a real-time, buffer-based algorithm. The ECG signal was continuously streamed, with each incoming data point stored in a dynamic buffer of fixed length. After the buffer was filled, the system determined the maximum amplitude of the signal and set a dynamic threshold at two-thirds of this maximum value (calculated as maximum amplitude - (maximum amplitude / 3). Any incoming sample exceeding this threshold was classified as an R-peak, allowing for adaptive detection based on real-time fluctuations in signal amplitude.

Instead of recording an explicit resting baseline period, the system automatically adjusted the R-peak detection threshold during the initial seconds of data acquisition. This brief adaptation phase allowed the algorithm to account for individual variations in heart rate and establish appropriate detection thresholds without the need for predefined calibration. Upon successful detection of an R-peak, the system immediately triggered a corresponding visual feedback response, ensuring precise alignment between participants’ physiological signals and the task’s visual stimuli.

### Psychological questionnaires

Prior to the heartbeat counting task, participants completed a battery of psychological questionnaires assessing anxiety, depression, body image, body and mental awareness. These factors can influence the perception of interoceptive signals and the response to emotionally charged stimuli. A brief description of the questionnaires used is given below. Full text of the questionnaires can be found in Supplementary materials.

1. *Multilevel Assessment of Interoceptive Awareness, MAIA-R*. The Russian adaptation of the MAIA-R Multilevel Assessment of Interoceptive Awareness MAIA-R ^59^ by Popova and Lopukhova ^72^ was used to assess interoceptive awareness. The 32-item instrument uses a 6-level Likert scale response format (0 = never, 5 = always;). In the Russian version, Cronbach’s alpha reliability coefficient, calculated for each of the scales, ranges from 0.6 to 0.85.
2. *The Body Image Questionnaire* by Skugarevsky and Sivukha was used to assess the level of dissatisfaction with one’s own body ^73^. Respondents are asked to rate 16 items on a 4-point scale (from 0 for “never” to 3 for “always”). In the article dedicated to the development of this tool, the authors provide a threshold value of 13 points, which may indicate disordered eating behavior and is recommended for use in screening studies, as well as an evaluation of the risk at 32 points, which indicates a high level of body dissatisfaction and a high risk of eating disorder in the respondent.
3. *The Hamilton Anxiety Rating Scale (HAMA)* quantifies the severity of anxiety and is often used to evaluate antipsychotic medications ^74^. It consists of 14 indicators, each of which is defined by a number of symptoms. Each indicator is rated on a 5-point scale from 0 (absent) to 4 (severe). The sum of all items provides a total score, with higher scores indicating greater severity of anxiety. Cut-off scores for the HAMA are as follows: mild Anxiety: total score of 14-17, moderate Anxiety: total score of 18-24, severe Anxiety: total score of 25 and above. Its internal consistency (Cronbach’s alpha) ranged from 0.77 to 0.92.
4. *Mindful Attention Awareness Scale (MAAS)*. A Russian adaptation of Mindful Attention Awareness Scale ^75^ by Golubyev ^76^ was performed to measure mindfulness. MAAS is a 15-item self-report tool designed to measure mindfulness, which is defined as a process of increased receptive awareness and attention to the present moment. Each item is rated on a six-point Likert scale from 1 (“almost always”) to 6 (“almost never”) and then added together to create a total sum score. Thus, lower scores indicate greater mindlessness, whereas higher scores indicate greater mindfulness. In previous studies, Cronbach’s alpha ranged from 0.80 to 0.90, indicating strong reliability.
5. *Emotionality questionnaire*. This diagnostic of emotionality was proposed by Suvorova in 1976 and determines the general emotionality of a person ^77^. The methodology includes 15 questions (statements). A score of 1 is assigned to each affirmative answer. The total sum of points is calculated. The internal consistency of the items in our sample is Cronbach’s α = 0.75. Interpretation: 0–5 points indicate low emotionality, 6–10 points indicate medium emotionality, and 11 or more points indicate high emotionality.
6. *Beck’s Depression Inventory*. This instrument is a 21-item, self-report questionnaire designed to assess and evaluate the frequency of anxiety symptoms over a one-week period ^78^. This test assesses two factors: cognitive and somatic symptoms. The instrument has good internal consistency (α = 0.92), test-retest reliability (r = 0.75; df = 81, P = <.001), and convergent and discriminant validity.

### Data analysis

Statistical analysis was conducted using Python scripts and the Statistica 64 software. Initially, to assess baseline differences between the experimental and control groups before the intervention, we used the chi-square test of independence for categorical variables and the Mann-Whitney U test for ordinal variables.

Next, we conducted a Mixed Linear Model Regression using the statsmodels.formula.api.mixedlm function in Python to evaluate changes over time, incorporating participant ID as a random factor to account for individual variability. The analysis revealed significant group-by-time interactions, indicating that the observed changes were driven by the training intervention. Subsequent correlation analysis was performed exclusively on variables that exhibited significant group differences (p-value < 0.05).

After, to evaluate within-group differences over time, we conducted the Wilcoxon signed-rank test, a non-parametric method suitable for paired comparisons that does not assume normality in the data distribution. Only variables demonstrating significant group differences (p-value < 0.05) were included in the subsequent correlation analysis.

Finally, to assess the relationships between changes in different variables over time, we applied Spearman’s rank correlation. Correlation analyses were conducted only for variables that showed statistically significant changes according to the Wilcoxon matched pairs test. Significant associations were identified at a threshold of p-value < 0.05. Relative changes in variables across time points were computed using the scipy library in Python. The relative change for each variable was calculated as:

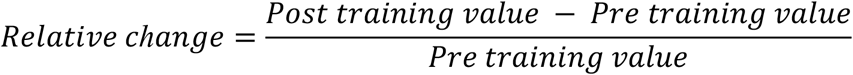

### Use of artificial intelligence tools

In compliance with the editorial policies of Scientific Reports, we disclose that Large Language Models (LLMs), such as ChatGPT, were utilized in the preparation of this manuscript. Specifically, these AI tools assisted in refining the language and enhancing the readability of the text. The authors maintained full control over the scientific content and interpretations presented. The use of LLMs was limited to editorial purposes and did not influence the scientific integrity of the work.

## Supporting information

Questionnaires

## Author contributions

Anna Rusinova and Maria Volodina: Contributed to conception and design. Anna Rusinova, Ekaterina Potapkina, Elizaveta Kozhanova, Vladislav Akimov: Contributed to acquisition of data. Vladislav Aksiotis: Assisted in developing the program code. Alexei Ossadtchi: Edited the article and acquired funding. Maria Volodina, Anna Rusinova: Contributed to interpretation of data. Anna Rusinova, Maria Volodina: Writing original draft, review, and editing.

## Acknowledgments

The authors would like to thank the participants for taking part in this research.

## Data availability statement

The data that support the findings of this study are available from the corresponding author upon reasonable request.

