## Supplementary material for "Interoceptive training enhances emotional awareness and body image perception: evidence from improved heartbeat detection and psychological outcomes": Questionnaires

### 1. Multidimensional Assessment of Interoceptive Awareness, MAIA-R

*Multilevel Assessment of Interoceptive Awareness, MAIA-R.* The Russian adaptation of the MAIA-R Multilevel Assessment of Interoceptive Awareness MAIA-R [1] by Popova and Lopukhova [2] was used to measure interoceptive awareness. The 32-item instrument uses a 6-level Likert scale response format (0 = never, 5 = always;). Cronbach's alpha reliability coefficient, calculated for each of the scales, ranges from 0.58 to 0.85. MAIA scales:

- Noticing: This factor assesses the extent to which an individual is conscious of physical signals within their body and can recognize subtle changes in bodily states.
- Not-Distracting: It reflects the ability to stay present with bodily sensations, even when they are uncomfortable, without trying to divert attention elsewhere.
- Not-Worrying: This factor evaluates how well an individual can experience unpleasant bodily sensations without adding a layer of worry or anxiety, contributing to overall emotional stability.
- Emotional Awareness: Assesses the ability to attribute specific physical sensations to physiological manifestations of emotions.
- Self-Regulation: Refers to the ability to regulate one's emotional state through attention to bodily signals.
- Body-Listening: Represents the active engagement in listening to the body for insights and guidance.
- Trusting: Relates to the belief that one's body is safe and reliable.

Instruction: Below is a collection of statements about your everyday experience. Using the 0-5 scale below, please indicate how frequently or infrequently you currently have each experience. Please answer according to what really reflects your experience rather than what you think your experience should be. Please treat each item separately from every other item.

1) When I experience tension, I notice where in the body that tension is located.
2) I notice states of bodily discomfort.
3) I notice which part of the body I am comfortable in.
4) I notice changes in my breathing, such as when it becomes faster or slower.
5) I don't notice (ignore) physical tension or discomfort until it increases.
6) I try to distract myself from the sensations of discomfort.
7) I try to overcome the sensation of pain or discomfort.
8) I get upset when I experience physical pain.
9) At any sensation of discomfort, I become anxious that something is wrong.
10) I am good at noticing unpleasant bodily sensations and not worrying about it.
11) I am able to pay attention to my own breathing without being distracted by what is going on around me.
12) I can maintain awareness of my inner bodily sensations even when there is a lot going on around me.
13) I keep track of my body posture when talking to another person.
14) I am able to return to awareness of my own body if I get distracted.
15) I can shift my attention from thoughts to bodily sensations.
16) I am able to maintain whole body awareness even when I experience pain or discomfort in a part of my body.
17) I am able to consciously focus attention entirely on my body.
18) I notice changes in my body when I am angry.
19) When something unpleasant happens in my life, I can feel it in my body.
20) I notice that in a calm situation my bodily sensations change.
21) I notice that my breathing becomes free and easy when I am comfortable.

22) I notice how my bodily sensations change when I am happy or joyful.
23) When I have an excess of strong experiences, I can find a quiet corner within myself. 24) When I am more aware of my body, I feel calmer.
25) I am able to control my breathing to release tension.
26) When I am overwhelmed by thoughts, I am able to calm myself by focusing on my body/breathing.
27) I am able to extract information about my emotional state from my bodily sensations. 28) When I am upset, I take my time to explore my bodily sensations.
29) I listen to my body to figure out what to do.
30) I feel comfortable in my body, like I am at home in my house.
31) I feel safe in my body.
32) I trust my bodily sensations.

Scores are between 0 and 5, where higher score equates to more awareness of bodily sensation. A percentile is also calculated, indicating how the respondent scored in comparison to a normative sample. Interpretation using percentiles helps contextualize scores. For example, percentiles below 50 indicate that the individual scored below what is typical. Extreme percentile scores (below 10 or above 90) are of particular clinical significance.

### 2. The Body Image Questionnaire

*The Body Image Questionnaire* by Skugarevsky and Sivukha was used to assess the level of dissatisfaction with one's own body [3]. Respondents are asked to rate 16 items on a 4-point scale (from 0 for "never" to 3 for "always"). In the article dedicated to the development of this tool, the authors provide a threshold value of 13 points, which may indicate disordered eating behavior and is recommended for use in screening studies, as well as an evaluation of the risk at 32 points, which indicates a high level of body dissatisfaction and a high risk of eating disorder in the respondent.
Instructions: Rate each statement on a four-point scale (0 - "never", 1 - "sometimes", 2 - "often", 3 - "always")

1. I don't like looking at myself in the mirror
2. Buying clothes draws my attention to how I look and is therefore unpleasant
3. I don't like to be noticed by others.
4. I avoid situations where others can see my body (e.g., going to the pool, beach, etc.).
5. I feel ashamed of my body in the presence of certain people.
6. I do not like my body
7. I feel that other people must think my body is ugly.
8. I feel embarrassed when friends and family members look at me.
9. I compare my body to others to see if they are fuller than I am.
10. I find it difficult to enjoy my activities because I feel embarrassed about my appearance.
11. I have feelings of guilt about my weight
12. I have negative thoughts and am self-critical/self-judgemental about my body and the way I look
13. I find it difficult to accept compliments about the way I look
14. When I look in the mirror, my attention is predominantly focused on parts of my body that need improvement
15. I feel humiliated and/or depressed in the presence of someone who I think is more attractive than I am.
16. I worry about my weight

One total score is calculated from the scale. To calculate raw scores, all scores on all items in the scale must be summed. A raw score value of 13 and above indicates a pronounced dissatisfaction with one's own body. The maximum score on the scale is 48.

#### 3. Hamilton Anxiety Rating Scale (HAM-A)

*The Hamilton Anxiety Rating Scale (HAMA)* quantifies the severity of anxiety and is often used to evaluate antipsychotic medications [4]. It consists of 14 indicators, each of which is defined by a number of symptoms. Each indicator is rated on a 5-point scale from 0 (absent) to 4 (severe).
Instruction: This questionnaire contains groups of statements. Read each group of statements carefully. Then identify one statement in each group that best describes how you have been feeling this week and today. For each

item, choose the value that best matches the severity of your symptoms. 0 - none, 1 - mildly, 2 - moderate, 3 - severe, 4 - very severe.

1. Anxious mood (preoccupation, expecting the worst, anxious fears, irritability)
2. Tension (feeling tense, startle, easily occurring tearfulness, trembling, feeling restless, inability to relax)
3. Fears (of the dark, strangers, loneliness, animals, crowds, transport)
4. Insomnia (difficulty falling asleep, intermittent sleep without rest, feeling broken and weak on waking, nightmares)
5. Intellectual impairment (difficulty concentrating, memory impairment)
6. Depressed mood (loss of usual interests, sense of enjoyment of hobbies, depressed mood, early awakenings, diurnal mood swings)
7. Somatic muscular symptoms (aches, twitches, tension, clonic convulsions, teeth grinding, voice breaking, increased muscle tone)
8. Somatic sensory symptoms (ringing in the ears, blurred vision, hot and cold flushes, sensations of weakness, tingling)
9. Cardiovascular symptoms (tachycardia, palpitations, chest pain, pulsation in blood vessels, frequent sighing)
10. Respiratory symptoms (pressure and tightness in the chest, choking, frequent sighs)
11. Gastrointestinal symptoms (difficulty swallowing, flatulence, abdominal pain, heartburn, feeling of a full stomach, nausea, vomiting, stomach rumbling, diarrhea, constipation, decreased body weight)
12. Genitourinary symptoms (rapid urination, strong urge to urinate, amenorrhoea, menorrhagia, frigidity, premature ejaculation, loss of libido, impotence)
13. Autonomic symptoms (dry mouth, red or pale skin, sweating, tension headaches)
14. Behaviour (fidgeting in chair, restless gesticulation and gait, tremors, facial frowning, tense facial expression, sighing or rapid breathing, frequent swallowing of saliva)

To obtain a total score reflecting the level of severity of the anxiety disorder, the scores for all items should be added together. In addition, the first six items can be evaluated separately as manifestations of anxiety in the mental sphere, and the remaining eight as manifestations of anxiety in the somatic sphere. Values of 17 points or less indicate the absence of anxiety, 18-24 points - the average severity of anxiety disorder, 25 points and above - severe anxiety.  The value of scores on all 14 items is ranked from 0 to 4. The total score takes values in the range from 0 to 56.

### 4. Mindful Attention Awareness Scale (MAAS)

*Mindful Attention Awareness Scale (MAAS).* A Russian adaptation of Mindful Attention Awareness Scale [5] was performed to measure mindfulness. MAAS is a 15-item self-report tool designed to measure mindfulness, which is defined as a process of increased receptive awareness and attention to the present moment. Each item is rated on a six-point Likert scale from 1 (“almost always”) to 6 (“almost never”) and then added together to create a total sum score. Thus, lower scores indicate greater mindlessness, whereas higher scores indicate greater mindfulness.

Instruction: Using a scale from 1 to 6, you were asked to rate how often or rarely something like this happens to you. The scale is inverted, i.e. the values with the highest frequency are on the left and the values with the lowest frequency are on the right. (1 point = Almost always, 2 points = Very often, 3 points = More often, 4 points = Rarely, 5 points = Very rarely, 6 points = Almost never).

1. I can experience an emotion and only realise it after a certain amount of time.
2. I drop or break things carelessly or inattentively, or because my mind is on something else. 3. I find it difficult to focus on what is happening in the present moment.
4. I have a tendency to go somewhere quickly without giving myself credit for what is going on around me or what happens along the way.
5. I don't notice signs of physical tension or discomfort until they become pronounced.
6. I almost always forget the other person's name when I am told it when I first meet them.
7. I often act in an automatic manner without realising what I am actually doing.
8. I practice most of my lessons without really concentrating.
9. I get so focused on goals that I lose contact with what I am currently doing to achieve them.
10. I do my work automatically without deeply focusing on it.
11. I happen to listen to people half-heartedly while doing something else.
12. I sometimes find myself in different places without knowing why or how I got there.
13. I worry about the future or the past.
14. Sometimes I do something without realising what or why I am doing it.
15. Sometimes I eat without realising that I am consuming food.

Assessment of results: The degree of your mindfulness in everyday life is assessed. To score, sum the answers for each item. The higher the sum, the more aware a person is in everyday life.

### 5. Emotionality questionnaire

*Emotionality questionnaire.* This diagnostic of emotionality was proposed by Suvorova in 1976 and determines the general emotionality of a person [6]. The methodology includes 15 questions (statements). A score of 1 is assigned to each affirmative answer. The total sum of points is calculated.

Instruction: Answer the test questions. The answer options are yes or no.

1. Can you blush so red from embarrassment or shame that you feel your cheeks flaming and tears welling up in your eyes?
2. Have you ever turned pale with fear or distress?
3. Are you often embarrassed, or are you shy?
4. Do you cry easily from resentment, unhappiness, empathy or even joy? Can you cry from aesthetic pleasure when you listen to music, read poetry?
5. Did sweat break out in an unpleasant or difficult situation?
6. Do you have a dry mouth when you are very excited? Does it make your voice sound dry? 7. In moments of great excitement or embarrassment, do you experience stiffness of the limbs, when your legs become stiff, "stilting" or "cotton" and shake?
8. Do you notice your fingers trembling when you are very excited or embarrassed, or do you have an internal shivering and chill-like state ("chills")?
9. Do you really get so excited before every performance that you feel like you have forgotten everything?
10. Can you lose your train of thought, get confused and stop talking while answering an exam or giving a public speech?

11 Do you often get angry and resentful? Can you get angry with a child and punish him/her in a hurry?

12. Do you tend to quarrel with your relatives when you see the injustice of their actions? Does it often end with

your tears, despondency and remorse?
13. You really can not switch off from troubles and griefs, not to think about them and bad mood completely owns you for a long time?
14. In moments of excitement or embarrassment you become excessively fidgety?
15. When you are excited, do you have pain in the solar plexus area?

For each affirmative answer ("Yes") 1 point is awarded. Interpretation of test results The more points a respondent scores, the higher his/her emotionality. - From 0 to 5 points - emotionality is low, - from 6 to 10 points - average, - from 11 points and above - high.

### 6. Beck's Depression Inventory

*Beck's Depression Inventory.* This instrument is a 21-item, self-report questionnaire designed to assess and evaluate the frequency of anxiety symptoms over a one-week period [7]. This test assesses two factors: cognitive and somatic symptoms. The instrument has good internal consistency (α = 0.92), test-retest reliability (r = 0.75; df = 81, P = <.001), and convergent and discriminant validity.
Instruction: The scoring scale is at the end of the questionnaire.

1.
0 I do not feel sad.
1 I feel sad
2 I am sad all the time and I can't snap out of it.
3 I am so sad and unhappy that I can't stand it.

2.
0 I am not particularly discouraged about the future.
1 I feel discouraged about the future.
2 I feel I have nothing to look forward to.
3 I feel the future is hopeless and that things cannot improve.

3.
0 I do not feel like a failure.
1 I feel I have failed more than the average person.
2 As I look back on my life, all I can see is a lot of failures.
3 I feel I am a complete failure as a person.

4.
0 I get as much satisfaction out of things as I used to.
1 I don't enjoy things the way I used to.
2 I don't get real satisfaction out of anything anymore.
3 I am dissatisfied or bored with everything.

5.
0 I don't feel particularly guilty
1 I feel guilty a good part of the time.
2 I feel quite guilty most of the time.
3 I feel guilty all of the time.

6.
0 I don't feel I am being punished.
1 I feel I may be punished.
2 I expect to be punished.
3 I feel I am being punished.

7.
0 I don't feel disappointed in myself.
1 I am disappointed in myself.
2 I am disgusted with myself.
3 I hate myself.

8.
0 I don't feel I am any worse than anybody else.
1 I am critical of myself for my weaknesses or mistakes.
2 I blame myself all the time for my faults.
3 I blame myself for everything bad that happens.

9.
0 I don't have any thoughts of killing myself.
1 I have thoughts of killing myself, but I would not carry them out.
2 I would like to kill myself.
3 I would kill myself if I had the chance.

10.
0 I don't cry any more than usual.
1 I cry more now than I used to.
2 I cry all the time now.
3 I used to be able to cry, but now I can't cry even though I want to.

11.
0 I am no more irritated by things than I ever was.
1 I am slightly more irritated now than usual.
2 I am quite annoyed or irritated a good deal of the time.
3 I feel irritated all the time.

12.
0 I have not lost interest in other people.
1 I am less interested in other people than I used to be.
2 I have lost most of my interest in other people.
3 I have lost all of my interest in other people.

13.
0 I make decisions about as well as I ever could.
1 I put off making decisions more than I used to.
2 I have greater difficulty in making decisions more than I used to.
3 I can't make decisions at all anymore.

14.
0 I don't feel that I look any worse than I used to.
1 I am worried that I am looking old or unattractive.
2 I feel there are permanent changes in my appearance that make me look unattractive
3 I believe that I look ugly.

15.
0 I can work about as well as before.
1 It takes an extra effort to get started at doing something.
2 I have to push myself very hard to do anything.
3 I can't do any work at all.

16.
0 I can sleep as well as usual.
1 I don't sleep as well as I used to.
2 I wake up 1-2 hours earlier than usual and find it hard to get back to sleep.
3 I wake up several hours earlier than I used to and cannot get back to sleep.

17.
0 I don't get more tired than usual.
1 I get tired more easily than I used to.
2 I get tired from doing almost anything.
3 I am too tired to do anything.

18.
0 My appetite is no worse than usual.
1 My appetite is not as good as it used to be.
2 My appetite is much worse now.
3 I have no appetite at all anymore.

19.
0 I haven't lost much weight, if any, lately.
1 I have lost more than five pounds.
2 I have lost more than ten pounds.
3 I have lost more than fifteen pounds.

20.
0 I am no more worried about my health than usual.
1 I am worried about physical problems like aches, pains, upset stomach, or constipation.
2 I am very worried about physical problems and it's hard to think of much else.
3 I am so worried about my physical problems that I cannot think of anything else.

21.
0 I have not noticed any recent change in my interest in sex.
1 I am less interested in sex than I used to be.
2 I have almost no interest in sex.
3 I have lost interest in sex completely.

INTERPRETING THE BECK DEPRESSION INVENTORY

Now that you have completed the questionnaire, add up the score for each of the twenty-one questions by counting the number to the right of each question you marked. The highest possible total for the whole test would be sixty-three. This would mean you circled number three on all twenty-one questions. Since the lowest possible score for each question is zero, the lowest possible score for the test would be zero. This would mean you circles zero on each question. You can evaluate your depression according to the Table below.

Total Score____________________Levels of Depression
1-10____________________These ups and downs are considered normal 11-16___________________ Mild mood disturbance
17-20___________________Borderline clinical depression
21-30___________________Moderate depression
31-40___________________Severe depression
over 40__________________Extreme depression
